# Generic Framework for Quantifying the Influence of the Mitral Valve on Ventricular Blood Flow

**DOI:** 10.1101/2022.08.25.505223

**Authors:** Jan-Niklas Thiel, Ulrich Steinseifer, Michael Neidlin

## Abstract

Blood flow within the left ventricle provides important information regarding cardiac function in health and disease. The mitral valve strongly influences the formation of flow structures and there exist various approaches for the representation of the valve in numerical models of left ventricular blood flow. However, a systematic comparison of the various mitral valve models is missing, making a priori decisions considering the overall model’s context of use impossible. Within this study, a benchmark setup to compare the influence of mitral valve modeling strategies on intraventricular flow features was developed. Then, five mitral valve models of increasing complexity: no modeling, static wall, 2D and 3D porous medium with time-dependent porosity, and one-way Fluid-Structure-Interaction (FSI) were compared with each other. The flow features velocity, kinetic energy, transmitral pressure drop, vortex formation, flow asymmetry as well as computational cost and ease-of-implementation were evaluated. The one-way FSI approach provides the highest level of flow detail, which is accompanied by the highest numerical costs and challenges with the implementation. As an alternative, the porous medium approach with our new expansion including time-dependent porosity provides good results with up to 10% deviations in the flow features (except the transmitral pressure drop) in comparison to the FSI model and only a fraction (11%) of numerical costs. Taken together, our benchmark setup allows a quantitative comparison of various mitral valve modeling approaches and is provided to the scientific community for further testing and expansion..

## 1 INTRODUCTION

For the heart, it is known that fluid dynamics within the ventricle play an important role in many aspects of physiology. There exists a relationship between fluid dynamical features and cardiac behavior in health and disease ^1,2,3^. Some examples include the vortex formation time for the diagnosis and prognosis of left ventricular heart failure ^4^ or the kinetic energy as a marker for cardiac function in single ventricle circulation ^5^. These simulation derived markers have the potential to corroborate clinical decisions and ultimately improve treatment of cardiovascular diseases. The mitral valve (MV), the valve that separates the left atrium (LA) and left ventricle (LV), plays in the central role in the development of the intraventricular flow structures. It is characterized by a highly complex non-regular three-dimensional shape. The motion of the MV is further influenced by the chordae tendineae, which are connected to the papillary muscles of the LV. This makes computational models very challenging and leads to several modelling approaches within studies focussing on cardiac blood flow.

Several studies have investigated the influence of the MV on ventricular blood flow by applying Fluid-Structure-Interaction (FSI) to simplified 2D models of the LV, LA, and MV ^6, 7, 8, 9^. Considering realistic three-dimensional geometries of the MV and the LV is extremely difficult. For this reason, most current studies apply nodal displacements to simplified MV geometries and the mitral annulus (MA) ^10, 11, 12^. Canè et al. ^13^. first implemented a one-way FSI simulation using overset meshes, also called Chimera grids, with mitral valve kinetics and torsional motion of a patient-specific LV. The resulting two conformally connected meshes could only undergo a limited degree of deformation due to the absence of remeshing and limited ability of smoothing techniques to handle nodal displacements. Despite the rather small MV opening angles, it was shown that the MV enhanced the propagation of the transmitral jet and promoted ventricular washout. Bavo et al. ^14^ directly imposed the valve motion determined from segmented ultrasound images and highlighted the importance of the valve on vortex dynamics and apical flow. A similar model using cardiovascular magnetic resonance imaging was developed by Chnafa et al. ^15^

Diode type models, in which the MV has no leaflets and the MA plane immediately changes from impermeable to permeable in early diastole, yielded controversial results. Seo et al. ^16, 17^ compared a physiological MV model based on MV opening angles obtained from *in-vivo* measurements with a diode type model ^18, 19^ and ^20^ and indicate that these simplified models do not produce physiological flow characteristics. In contrast, Saber et al. ^21^ and Vedula et al. ^19^ were able to show good qualitative agreement with *in-vivo* experiments using the same simplified approach. This idea was further extended by implementing a planar MV model with variable orifice area, which is based on imaging techniques and mimics the physiological effective orifice area of the MV ^22, 23^ and ^24^. Daub et al. adapted this approach by thickening the MV plane and adding porous properties to the effective orifice area ^25^. By tilting the MV plane, they were able to reproduce the asymmetry of the transmitral jet. They compared this implementation with a physiologically inspired 3D valve with predefined motion within the same ventricular geometry. The ventricular walls were defined to move while the LA remained fixed. Their results revealed enhanced vortex formation and improved ventricular washout of the approach with porosity compared to classical diode type models.

Other groups such as Liao et al. ^26^ implemented an approximated MV using parametric equations, which was first presented by Domenichini and Pedrizzetti ^12^. The MV was defined as a static wall that changes abruptly from closed to open in response to the pressure in the LA. This led to jumps in the solution data and can generally cause numerical instabilities.

Inconsistent results of existing modeling approaches for the MV make it difficult to establish clear decision criteria regarding what type of MV model should be chosen, considering the model’s context of use. This uncertainty in the scientific community leads to a lot of trial and error in the design of computational models. The central questions that often arise are: How simple can the MV be modeled to still obtain reliable results? What is the error compared to a more realistic representation? How does it relate to the computational costs and ease-of-implementation?

The differences in the modeling setups in which the MV approaches are tested are particularly problematic, as each requires different simplifications and a variety of pre-processing steps. To obtain a better understanding of the quality of the different modeling approaches already presented in the literature, a simple framework will be created to make the assessment easier and more flexible. This way, the ventricle-valve interaction, which often interferes the validity of the obtained results, will be excluded.

Previous studies have focused mainly on a qualitative evaluation of vortex formation and propagation and therefore provide only a rather basic understanding of the valvular influence on flow features of ventricular blood flow. This makes it difficult to decide, which modeling approach is most suitable for specific applications in terms of computational efficiency and accuracy. However, it would be desirable to choose the level of detail of an implementation based on the flow information needed. For this reason, this study focuses on a quantitative comparison of widely used flow features using existing modeling approaches of the MV within a benchmark setup..

## 2 METHODS

At total, five different approaches of mitral valve modeling were compared with each other. One-way Fluid-Structure-Interaction (FSI), porous zone (PZ), porous jump (PJ), static wall (SW) and no valve modeling at all (NO). For all comparisons, the one-way FSI setup was taken as the ground truth and the outcomes of the other models were assessed with respect to the FSI simulation. To justify this assumption, a two-way FSI simulation with a heart valve inspired benchmark from Gil et al. ^27^, Hesch et al. ^28^ and Wick et al. ^29^ was performed and showed an excellent agreement with the existing results. The correct modeling of porosity was additionally confirmed with a simple analytical solution given for the pressure drop across a porous medium ^30^. For both preliminary studies see the Supplementary Material.

All simulations were performed with the ANSYS R2021 (ANSYS Inc., Canonsburg, US) software package. The flow simulations were performed with ANSYS Fluent and the structural simulations with ANSYS Mechanical. The coupling of these two modules was established with ANSYS System Coupling. The benchmark setups are available at 10.5281/zenodo.6741579.

### 2.1 Setup of benchmark environment

All the different modeling approaches were tested in a benchmark setup shown in figure 1. The setup consisted of a tube with a sudden expansion with circular and non-circular cross sections illustrated in AA and BB. The MV was placed exactly at this transition, where the dimensions of the fluid domain roughly double. In addition, the inlet and outlet regions, labeled as one and three in figure 1, were extended to ensure fully developed flow near the inlet and outlet boundaries. To save computation time, the flow domain was split into three regions with different mesh densities and element types.

**FIGURE 1.**
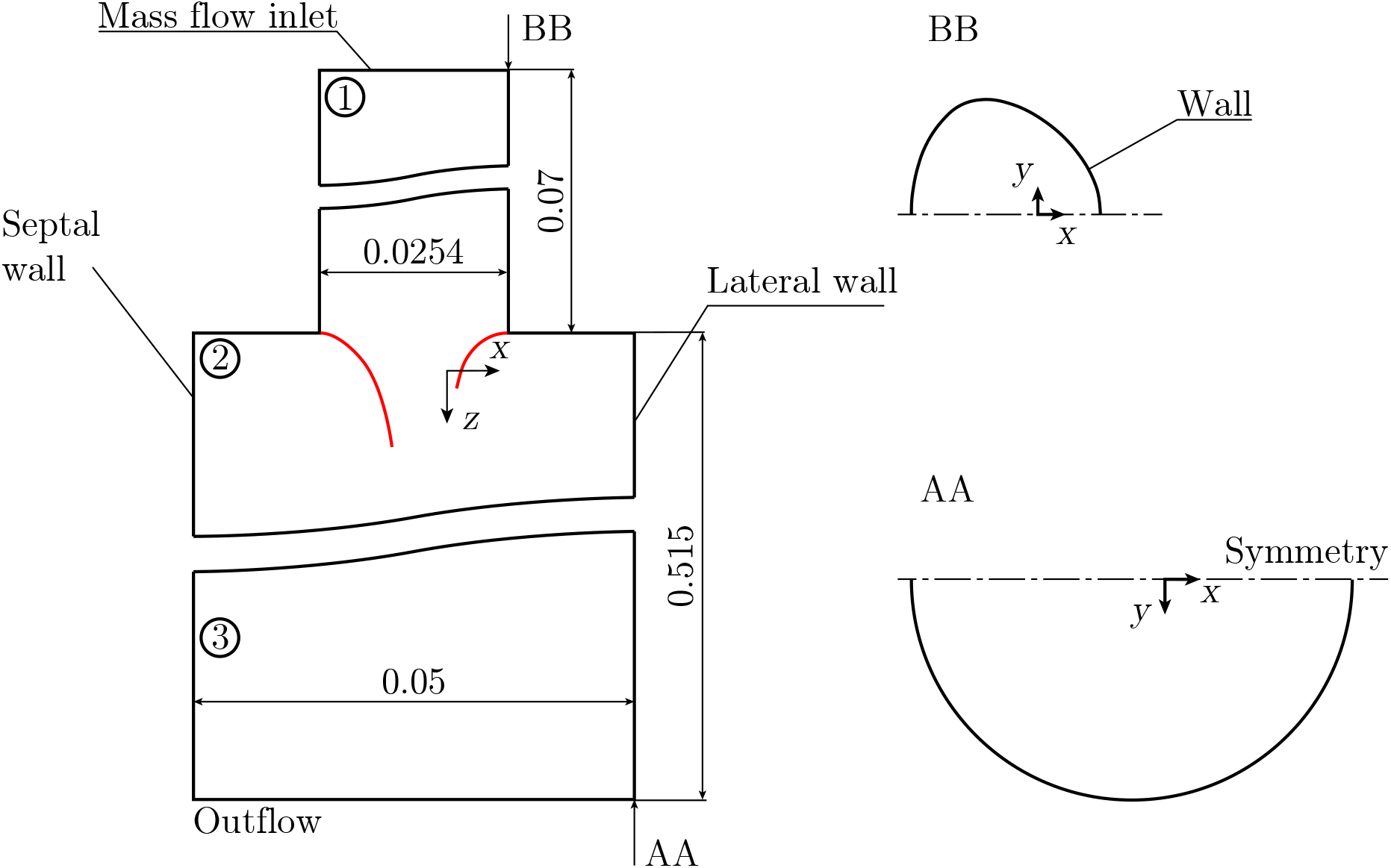
Geometry and boundary conditions of the heart valve benchmark setup. Dimensions in [m]. Mirror at y-axis!

The mass flow rate waveform shown in figure 2 was imposed at the inlet and had an E/A ratio of 1.35^31^ and mass flow rate values similar to the study by Seo et al. ^17^. A heart rate of 75 bpm was assumed with a diastole of 0.6 s and a systole of 0.2 s.

**FIGURE 2.**
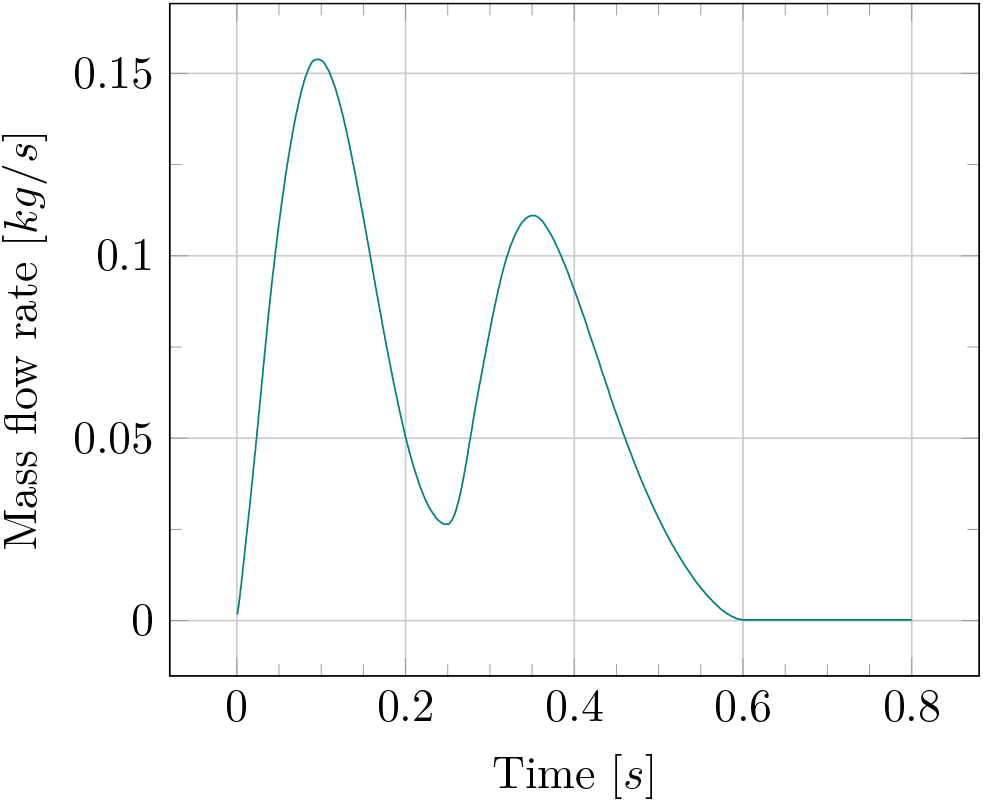
Flow rate wave form through MV with E/A ratio of 1.35.

The flow at the exit point was constrained by an outflow boundary condition, by which the diffusion flux was set to zero for all flow variables and an overall mass balance correction was applied. Further, the cylinder walls had no slip boundary conditions. The domain was split in half, since the problem is symmetric with respect to the *xz*-plane. Blood was modeled as an incompressible Newtonian fluid with a density of *ρ* = *μ* 1056.4 kg m^3^ and viscosity = 0.0036 Pa s and the flow was considered to be fully laminar. At total, three cycles were simulated.

Due to the subsonic incompressible flow field, a pressure based solver with a segregated solution scheme was used. The convergence criterion for each time step required that the locally scaled residuals decrease to 1 × 10^−5^ for each equation solved. In addition, convergence trends were documented for the area-averaged velocity magnitude and static pressure at the symmetry plane, which had to satisfy a convergence criterion of 1 × 10^−4^. A bounded second order implicit transient formulation was used for the time discretization. Furthermore, a second order upwind scheme for the convective terms was used. The time step size was chosen to be 5 × 10^−4^ s. The average flow Reynolds number was around 1756, and the Womersley number was 26, both of which were based on the hydraulic diameter of the mitral annulus (MA) and the heart rate. Both values combined gave a Strouhal number of 0.06 and are within the range for a healthy adult human heart ^32^.

### 2.2 Mitral valve modeling approaches

This section contains a detailed description of the different modeling approaches. First, the most complex implementation using one-way FSI coupling is explained and then simplification approaches are consecutively described

### One-way Fluid-Structure-Interaction

In order to perform a one-way FSI simulation, a 3D valve geometry needed to be implemented. The geometry is shown in figure 3. It was constructed in its closed position, had a D-shaped opening area and a non-circular annulus. The whole valve was assumed to have a uniform thickness of 6 × 10^−4^ m, see ^33^. A fixed support boundary condition was prescribed for the annulus ring. The stress-strain relation of the MV leaflets was described by a hyperelastic neo-Hookean material model with a density *ρ* = 1100 kg m^3^, the Poisson’s ratio *ν* = 0.488 and the Young’s modulus *E* = 4.1 × 10^6^ Pa and 2 × 10^6^ Pa for the anterior and the posterior leaflets, respectively ^34^. Opening and closing of the valve was achieved through a transient structural simulation requiring an opening angle of the leaflets of about 90° and an opening area in the range of physiological values, see figure 3, as well as a duration of the opening and closing process of 0.1 s. The applied boundary conditions on the MV are listed in the Supplementary Material.

**FIGURE 3.**
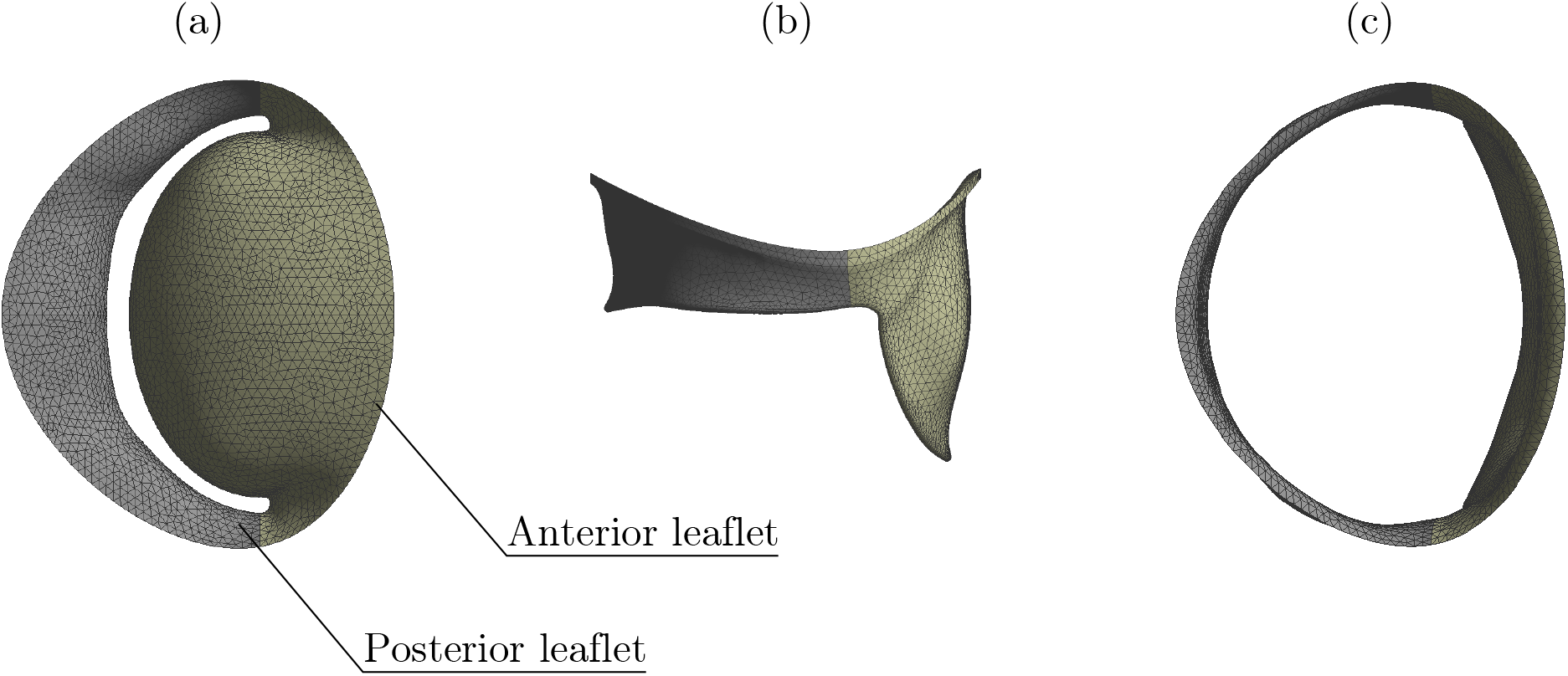
(a) Fully closed in axial view. (b) Fully open in coronal view. (c) Fully open in axial view.

The solid domain was meshed using tetrahedral elements of size 6 × 10^−4^ m, resulting in one mesh element per thickness of the valve. Since the valve was modeled as a nearly incompressible material, having a poisson’s ratio close to 0.5, the more robust mixed u-P formulation was used in the structural solver together with a nonlinear stabilization method which uses an energy dissipation ratio to consider convergence instabilities due to large displacements of the leaflets.

The fluid domain was meshed with tetrahedral elements and an element size of 0.001m in the main part of the benchmark setup. The extended inlet and outlet sections were meshed with an element size of 0.0025m. This resulted in a mesh with 1.38M elements and was used for all simulations. The according mesh independence study is included in the Supplementary Material. Since remeshing occured every time step and the second order in time transient formulation is generally incompatible with dynamic remeshing, the time advancement accuracy was reduced to first order. The SIMPLEC algorithm was used, because an increased under-relaxation can be applied to the pressure-correction. To increase convergence speed-up, the under-relaxation factor for the pressure correction was set to 1.0 and a skewness correction of 1 was used. The latter is particularly beneficial for cases with high mesh skewness. The number of inner iterations per time step was set to 20 in order to reach convergence of all relevant flow features for each time step. The mesh of the fluid domain deformed according to the opening and closing of the MV. In order to ensure a good mesh quality throughout the whole simulation, diffusion-based smoothing and both local cell and local face remeshing methods were used. The diffusion-based smoothing used the diffusion coefficient of *γ* = 1. The remeshing used the minimum edge length of *L*_min_ = 5 × 10^−4^ m and the maximum length *L* _max_ = 0.001/m at a remeshing interval of 1. The upper thresholds of cell and face skewness were set to 0.7 and 0.5, respectively.

A transient one-way FSI was performed by using the system coupling module with the surfaces in contact with the fluid defined as fluid solid interfaces. As there was only one way transfer between structural and fluid simulation, the number of system coupling iterations was set to one, which made it obsolete to specify any values for the RMS residuals.

### Orifice like model using porous media

In this approach, the MV was implemented using the laws of porous materials, as suggested by Daub et al. ^25^. By inserting a cell zone of porous medium into the MV plane, the transmitral flow was regulated. The geometric representation was obtained by projecting the physiological MV in its open and closed positions, as shown in figure 3, onto the MV plane. As a result, three porous zone faces were created with a fully permeable region, a fully impermeable region, as well as a region with a time-dependent resistance, which are illustrated in figure 4.

**FIGURE 4.**
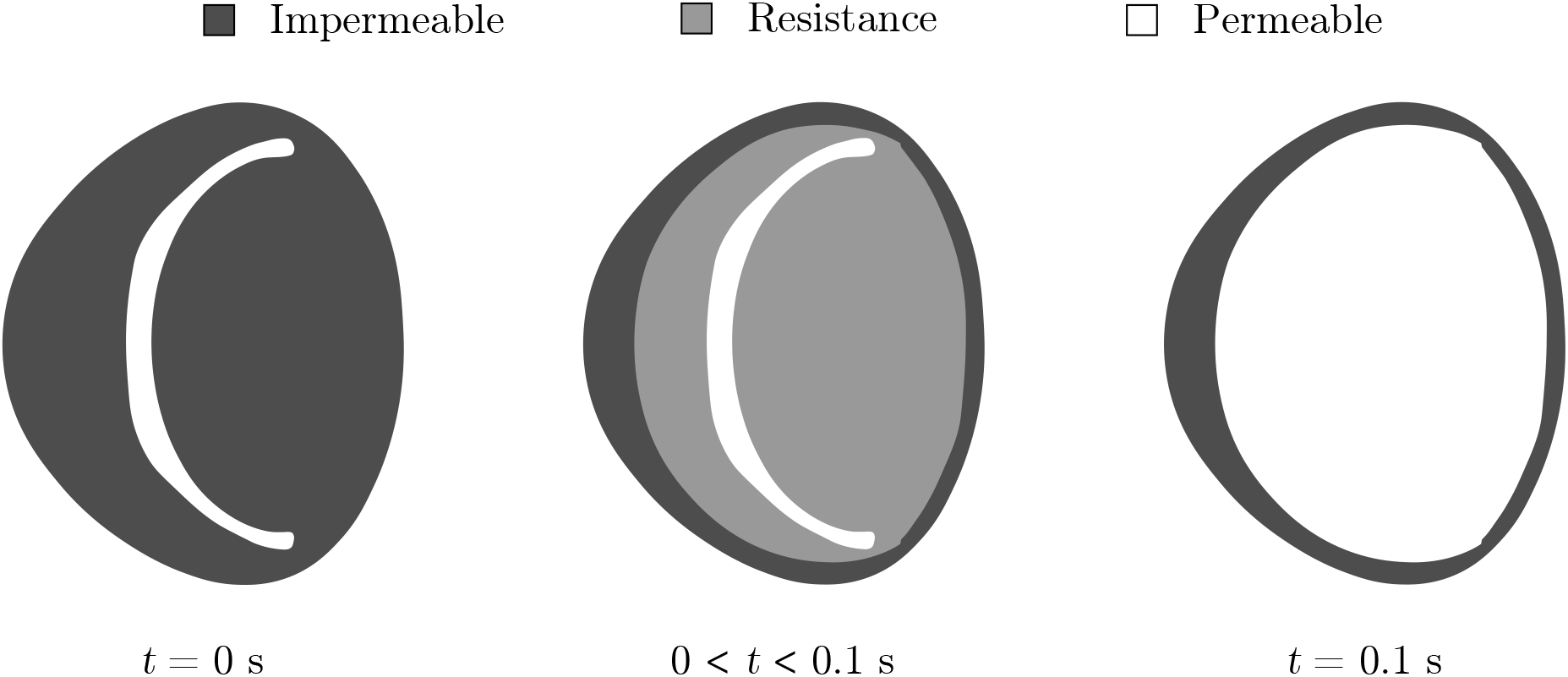
Projection of 3D MV onto inclined 2D plane. Three areas with different resistances during one cardiac cycle

To account for the asymmetry of the flow, the MV plane was inclined by 6°. Moreover, the resulting surfaces were thickened by 0.0012 m, giving the final assembly of the main fluid domain shown in Supplementary Figure 7, right side. A poly-hexcore mesh with 340 000 cells was created, using an element size of 0.001/m for the main section of the flow and 0.002 m for the inflow and outflow regions. The porous cell zones were discretized using an element size of 2.5 × 10^−4^ m.

The porosity of the cell zones was modeled using Darcy’s law, which is typically used for laminar flows with low Reynolds numbers. It relates the pressure drop across a porous medium to the velocity of the fluid. To additionally account for the dynamic head at high velocities, a correction for inertial losses, known as the Forchheimer term, was added. The resulting equation for the pressure drop in each direction reads

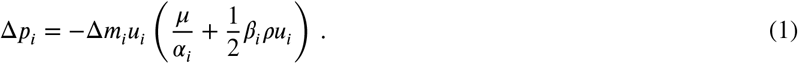

Here, *α*_*i*_ is the permeability of the porous zone in each direction, ranging from one, which corresponds to no resistance, to zero, denoting a high resistance. Further, *u*_*i*_ and Δ_*m*__*i*_ are the velocity components and the thicknesses of the medium in *x, y* and *z*-direction. The correction for inertial losses is provided by the Forchheimer coefficient *β*_*i*_, which usually has to be determined experimentally. Because the implementation is only intended to mimic the characteristic pattern of transmitral flow, both *α*_*i*_ and *β*_*i*_ were defined such that they block the flow in the tangential and bi-tangential directions of the MV plane. This was achieved by setting their values several orders of magnitude higher compared to those in the flow direction.

The inverse permeability *α*^−1^ and the Forchheimer coefficient *β* normal to the MV plane were defined as follows for the region with variable resistance

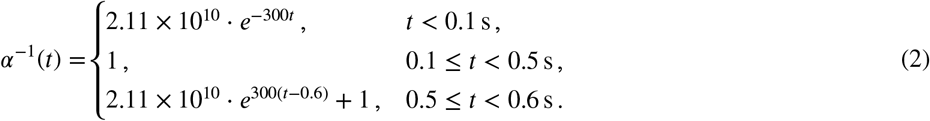

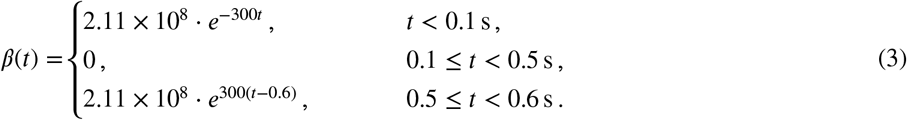

It can be seen that *α*^−1^ and *β* were set to 2.11 × 10^10^ 1/m^2^ and 2.11 × 10^8^ 1/m, respectively, when the MV is fully closed at *t* = 0 s. As the MV starts to open, the values drop exponentially to one and zero, resulting in no resistance for that period of time. As soon as the MV starts to close again, the values increase again to their initial value at *t* = 0 s. The profile for the whole diastole is shown on the left side of Supplementary Figure 7 and the implemented code within Fluent can be found in the Supplementary Material. The same type of functions was implemented for the other directions, but with values in the interval of 2.11 × 10^5^ < *α*^−1^ ≤ 2.11 × 10^10^ 1/m^2^ and 2.11 × 10^5^ < 2.11 × 10^8^ 1/m, respectively.

Again, SIMPLEC was used with a skewness correction of 1.0 and an under-relaxation factor for the pressure correction of 0.7. Additionally, the frozen flux formulation was employed, which enables to discretize the convective term of the generic transport equations using the mass flux at the cell faces from the previous time level. As generally recommended for 3D simulations on polyhedral meshes, which may not have perfectly planar faces, warped-face gradient correction (WFGC) was set. WFGC helps preventing accuracy degradation due to very high aspect ratios, cells with non-flat faces and highly deformed cells with high skewness. Further, high order term relaxation was used. Standard initialization and 50 iterations per timestep were used to reach convergence.

In an alternative approach, the 3D porous cell zones were reduced to the porous zone faces shown in figure 4. A porous jump boundary condition was then defined for the face with variable resistance, which is essentially a 1D simplification of the previously described porous zone model. Here, the pressure drop across the MV plane was also defined as a combination of Darcy’s law and the Forchheimer term. The face permeability *α* and the Forchheimer coefficient *β*, were defined for the boundary with variable resistance as follows

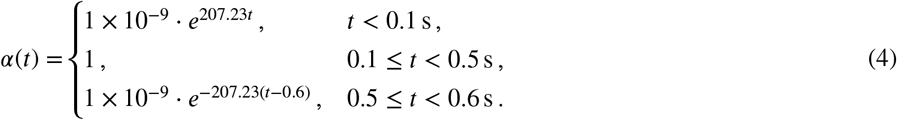

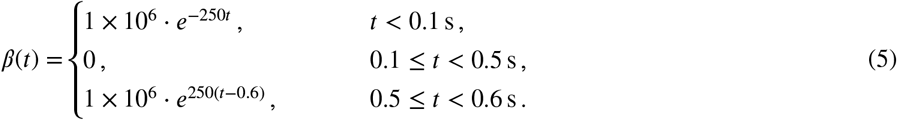

The implementation within Fluent using nested if statements in the expression editor can be found in the Supplementary Material. Further, the porous medium thickness was set to 0.0012 m. In contrast to the porous zone implementation, this approach used the SIMPLE scheme for the pressure-velocity coupling with an under-relaxation factor for the pressure correction of 0.3. All other simulation parameters remained unchanged.

The fluid domain was again discretized using a poly-hexcore mesh with 536 000 elements. Mesh parameters were generally identical to those of the porous zone model.

## Static wall

Another approach was implemented that modeled the MV as an internal baffle only. The 3D model of the physiological MV shown in figure 3 was reduced to a surface model of the MV in its fully open state. In order to consider that no flow passes the MV during ventricular systole, the benchmark setup was expanded by an additional rectangular outlet region as shown in Supplementary Figure 8 and time-dependent expressions for the flow rate weighting at the two outlets were defined. In consequence, for *t* < 0.1 s and *t* ≥ 0.5 s no flow passed through the MV and, when the MV was fully opened for 0.1 ≤ *t* < 0.5 s, all the flow went through the main outlet. The implemented expressions can be found in the Supplementary Material. The adapted flow domain was discretized using a poly-hexcore mesh with 270 000 elements. Different to all the other approaches, both the main part of the flow domain and the inlet region were defined to have an element size of 0.001/m. The segregated solver SIMPLEC was used with a skewness correction of 1.0 and an under-relaxation factor for the pressure correction of 0.7. Again, frozen flux formulation and WFGC were used. The flow field was solved using hybrid initialization and 20 iterations per time step.

### No modeling

The simplest of all approaches did not consider the MV at all, resulting in a simple pipe flow without any obstruction. A poly-hexcore mesh with 220 000 elements was used. Compared to the previously described approaches, the under-relaxation factor for the pressure correction was increased to 1.0, giving a higher rate of convergence. Moreover, only WFGC was used to stabilize the simulation.

### 2.3 Evaluation approaches

At total five different setups (FSI, PZ, PJ, SW, NO) were compared with each other and the following outcomes were of interest: axial velocity, kinetic energy density, diastolic pressure drop, vortex formation, jet propagation speed and the flow asymmetry *S*. The parameter *S* quantified the level of asymmetry and was based on the fact, that any function can be decomposed into a symmetric *u*_+_ and skew-symmetric part *u*_−_:

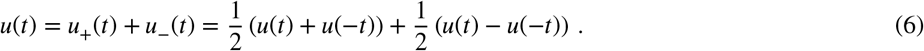

The asymmetry was then measured by applying the fol2lowing Euclidean n2orm

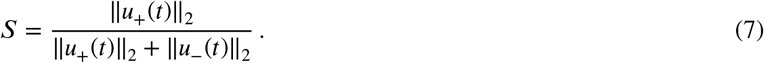

This was conducted for each time instant and then the average was taken over the whole diastole. This yielded a global metric that gives values from 0, fully symmetric flow, to 1, fully asymmetric flow. At last, the numerical costs and the ease of implementation were assessed.

## 3 RESULTS

### 3.1 Velocity, kinetic energy and pressure drop

The velocity in the *z*-direction at a reference point below the MV is evaluated for the diastole of each cycle in figure 5. The axial velocity of the different approaches follows a similar characteristic shape for each cycle. However, only in the remeshing case a cycle-independent solution can be obtained.

**FIGURE 5.**
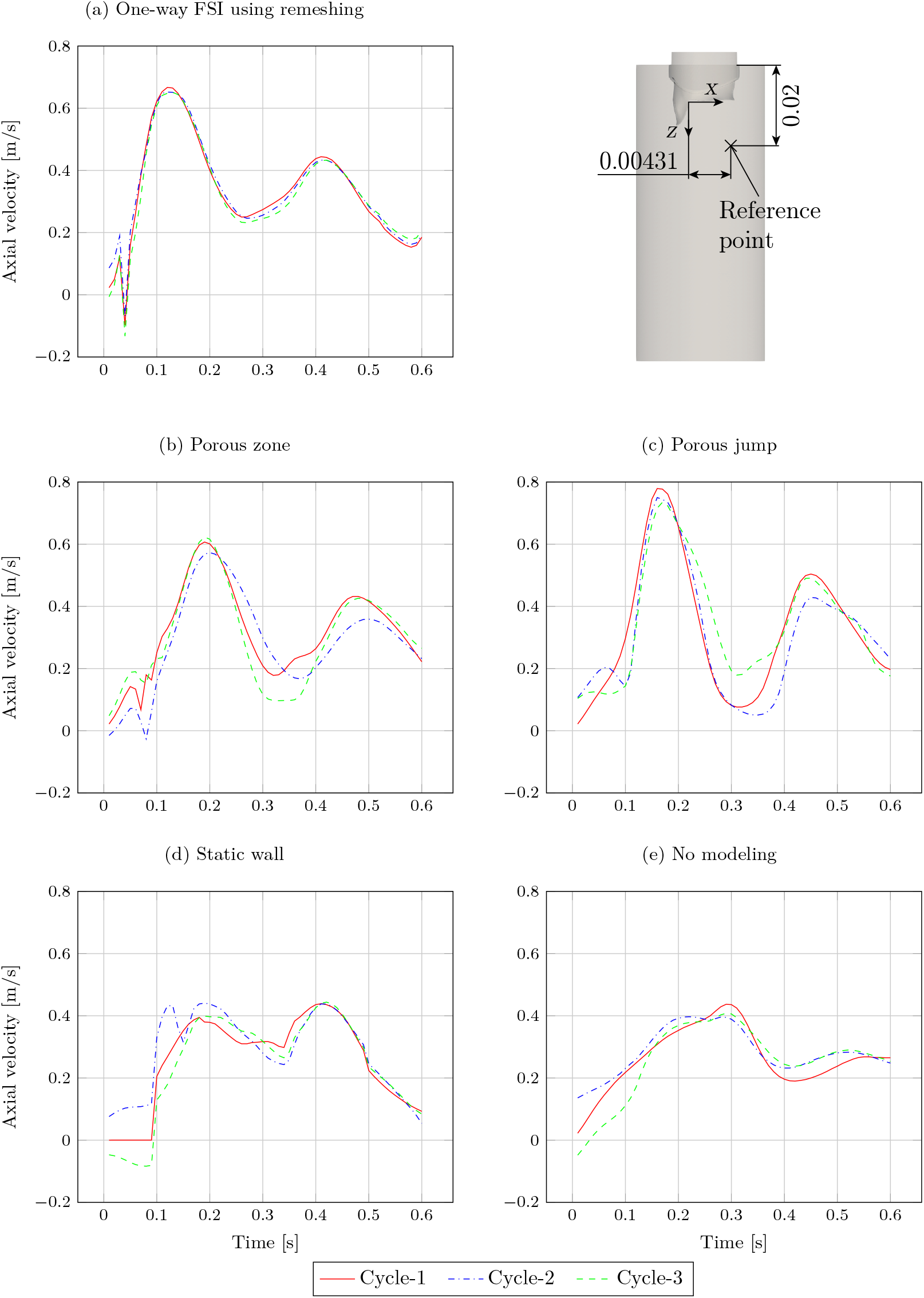
Axial velocity at a reference point below the MV for each cycle. Dimensions in [m].

For all cases, there exist phase shifts of the E- and A-waves as well as deviations of the maximal velocities when compared to the FSI case. In the porous zone case, there is a delay of about 0.07 s and 0.06 s, for the E- and A-wave and lower maximal velocities of 9% and 2%, respectively. The porous jump case exhibits a shift of about 0.04 s and increased peak velocities of 16% and 14%.

Comparing the phase shift of the simulation using porous zone to the simulation that uses a dynamic mesh, it can be observedhat the peak velocities of the E- and A-wave are shifted by about 0.07 s and 0.06 s, respectively. Both maxima of the simulation with a porous jump condition, however, are only delayed by about 0.04 s. In contrast, for the simulation with the MV being a static wall, only the E-wave is shifted by 0.06 s. The largest difference is found for the simulation without any modeling of the MV. Here, the E- and A-wave are delayed by about 0.17 s and 0.14 s, respectively. Comparing the peak velocities of the E- and A-wave of the porous zone model with the model using a dynamic mesh, it can be clearly seen that the model with porosity gives slightly lower values reduced by about 9% and 2%, respectively. In contrast, the maxima of the simulation using a porous jump condition are increased by about 16% and 14%. As already observed for the phase shift, the simulation using a static wall gives the same results for the A-wave as the simulation with remeshing. The E-wave, however, is about 40% slower. Once again, the simulation without an MV yields the largest deviation compared to the one using remeshing, which is about −34% and −39%, respectively.

To compare the shape of the velocity distribution of the simulation using remeshing to all other approaches, the Spearman rank correlation coefficient *r*_s_ was used. The highest correlation is found for the simulation with porous medium with *r*_s_ = 0.92, which can be qualitatively confirmed with figure 5. The simulation with a porous jump condition shows a comparably high similarity with *r*_s_ = 0.91. The correlation to the simulation which implements the MV as a static wall, however, is only *r*_s_ = 0.61. The simulation without modeling the MV shows the lowest correlation with *r*_s_ = 0.46. The p-values for all approaches were *<* 0.05.

The diastolic kinetic energy per unit volume (energy density) of the main fluid domain is visualized in figure 6. The energy density for the simulation with remeshing first increases suddenly and then drops slightly again before reaching its maximum at approximately *t* = 0.1 s. As the transmitral flow is zero for *t <* 0.1 s in the static wall modeling approach, the energy density is extremely low for this time interval. For *t* = 0.1 s the energy density increases stepwise. A similar behavior can be observed for *t* = 0.5 s. The energy density in the remeshing case is highest for the first peak of the inlet velocity profile. The model using a porous jump condition has a comparably high value. For all other approaches the energy density is lower. For the A-wave, however, the porous jump model overestimates the energy density and gives higher values than the one-way FSI model. In contrast, the energy density of the models using porous zone and the MV as static wall is comparable to the remeshing case for this time interval. The simulation without any modeling of the MV underestimates the energy density for the entire diastole. Comparing the average values of the diastolic energy density, one can observe that the simulation with porous zone provides a value that is about 11% lower than the one obtained with the remeshing model. In contrast, the simulation using a porous jump condition yields an average value that is about 10% higher. The simulations where the MV is just a static wall and the model without an MV produces results that are about 32% and 37% lower, respectively.

**FIGURE 6.**
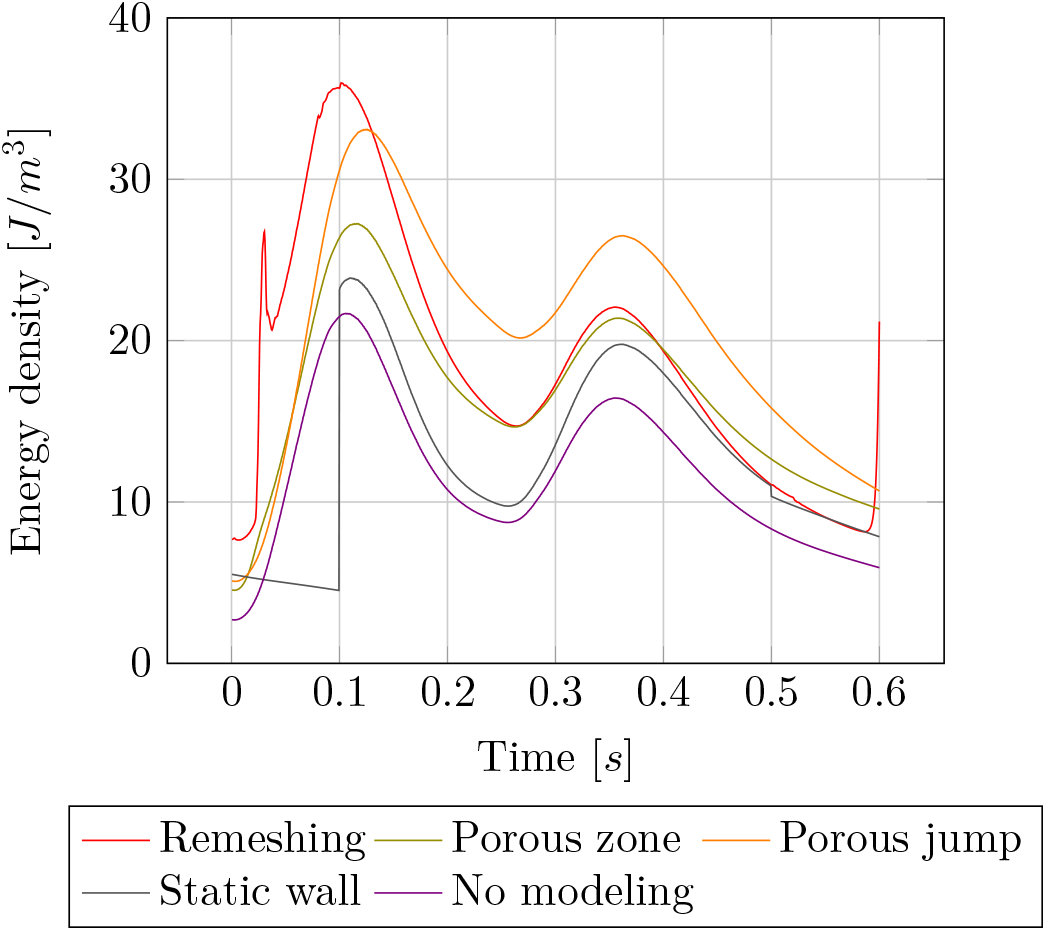
Energy density (kinetic energy per unit volume) in diastole of the third cardiac cycle

The averaged transmitral pressure gradients for the various modeling approaches are: 78*P a* for the remeshing model, 95*P a* for the model with a porous zone and 74 Pa with a porous jump, and 38*P a* and 33*P a* for the static wall and no MV model, respectively.

### 3.2 Vortex formation

Figure 7 and 8 show vortex structures during the filling phase of both E- and A-wave of the cardiac cycle by visualizing the isosurfaces of the Q-Criterion. To illustrate the direction of rotation, they are colored by the axial velocity, while a positive velocity in *z*-direction indicates towards the outlet of the fluid domain. They show the formation, pinch-off and propagation of both primary and secondary rings. For the remeshing model, which is shown in figure 7 (a), the propagation of the almost circular vortex ring, which is labeled as primary ring, at *t* = 0.1 s towards the lateral wall of the tube can be observed. As the vortex continues to propagate in the direction of flow, the primary ring begins to disintegrate at *t* = 0.2 s. At *t* = 0.3 s the disintegration is complete and the secondary ring begins to form at the leaflets of the MV. After pinching off, the second vortex ring continues to propagate further towards the outlet of the tube. As it can be seen in figure 7 (b) and 8 (b), both the simulation that models the MV as a porous zone and the simulation that considers it only as a static wall produce vortex structures that equally move towards the lateral wall of the tube. The other approaches are not able to reproduce this feature. Comparing the penetration depth of the vortices into the tube in general, it can be observed that the models with porous medium have a similar propagation speed as the one-way FSI remeshing simulation. In the other modeling approaches, the vortices move slower. Furthermore, the vortices are more stable and therefore start to disintegrate later in all other approaches, except the simulation without any representation of the MV. In the latter, the disintegration is already completed at *t* = 0.3 s, while in all the other simplified cases this process starts just at about *t* = 0.4 s.

**FIGURE 7.**
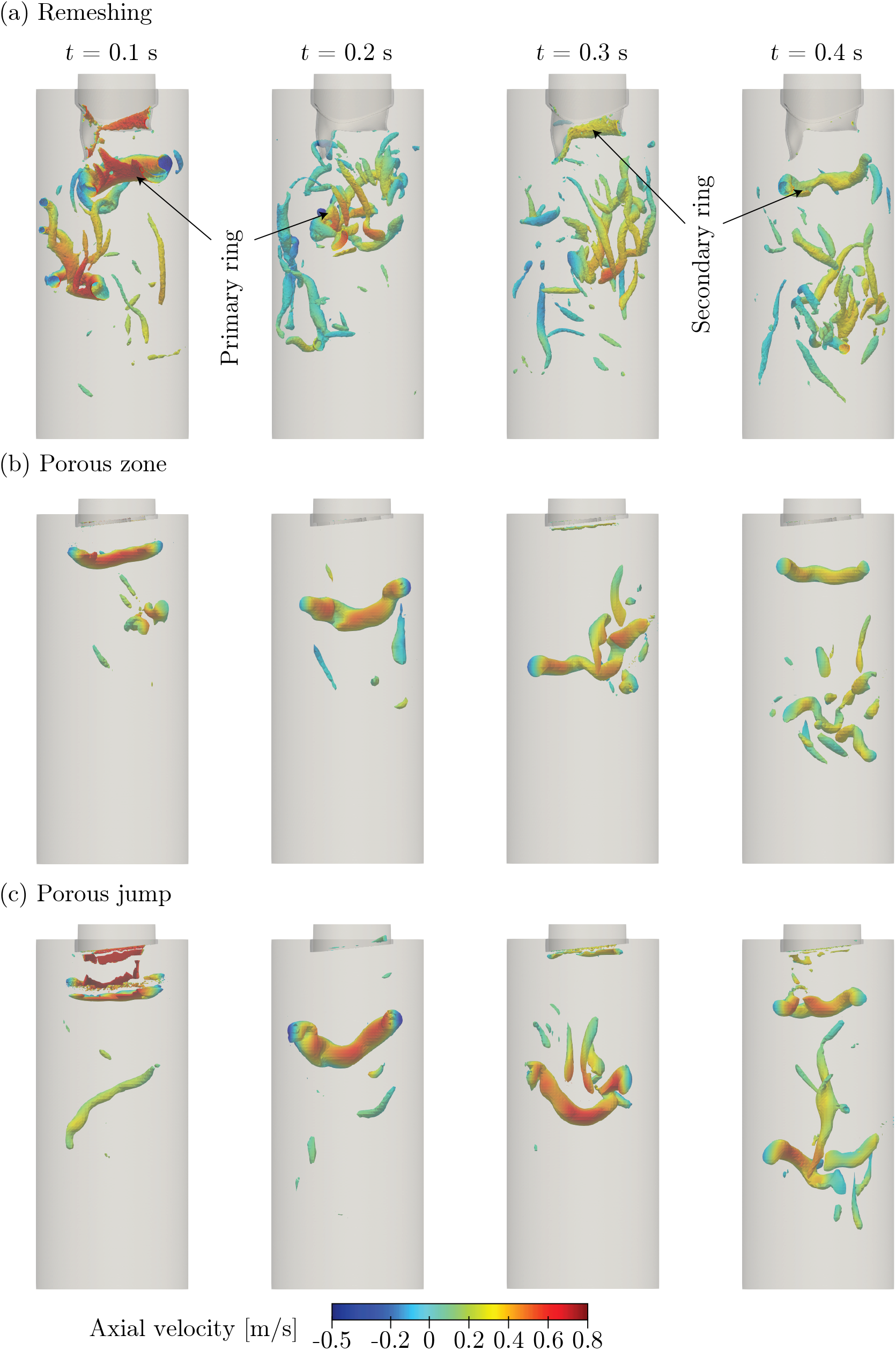
Vortical structures visualized by iso-surfaces of the Q-criterion with constant value defined to be 4000. Colored by axial velocity, with a positive value indicating main flow direction.Comparison of remeshing with porous zone and porous jump approaches.

**FIGURE 8.**
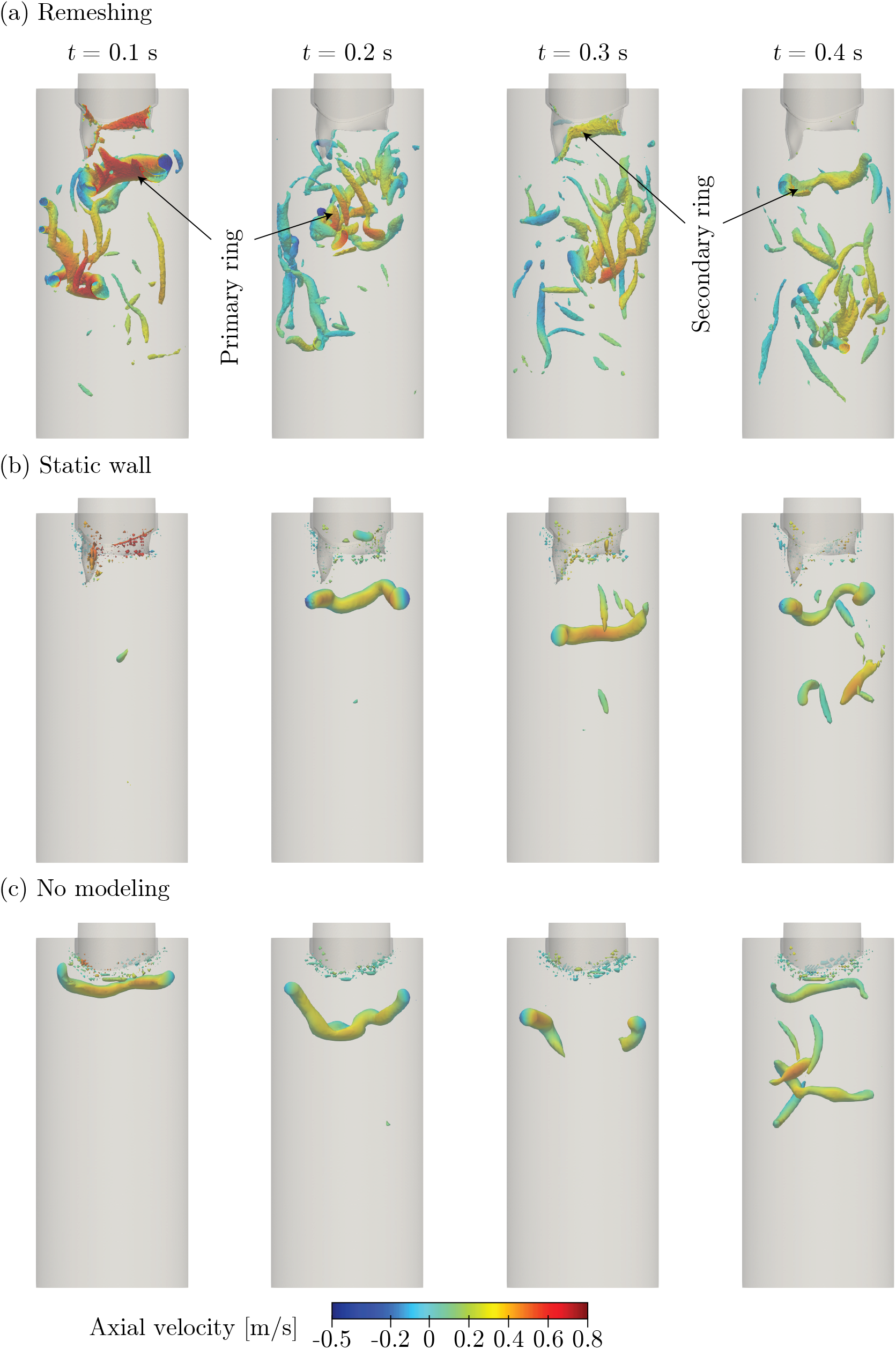
Vortical structures visualized by iso-surfaces of the Q-criterion with constant value defined to be 4000. Colored by axial velocity, with a positive value indicating main flow direction.Comparison of remeshing with static wall and no MV approaches.

### 3.3 Flow asymmetry

A key feature of the diastolic flow pattern in the left ventricle is the septal-lateral asymmetry of the flow velocity. The temporal change of the axial velocity was monitored along a line below the MV in the *x*-direction, which is indicated as *line-x* in figure 9. The peak velocity of the one-way FSI simulation shifts from being at around *x* = 0.03 m for the early filling phase (E-wave) at *t* = 0.025 s to the lateral wall at *x* = 0.015 m for *t* = 0.1 s, when the MV is completely open. The same behavior can be observed for the simulation using porous zone, yet the maximum axial velocity is shifted towards the septal wall and the incoming jet is delayed by about *t* = 0.05 s. The same holds true for the simulation, where the MV is only a static wall. In contrast, the overall flow pattern in the septal-lateral direction is quite symmetric for the approach with a porous jump condition and without any modeling of the MV.

**FIGURE 9.**
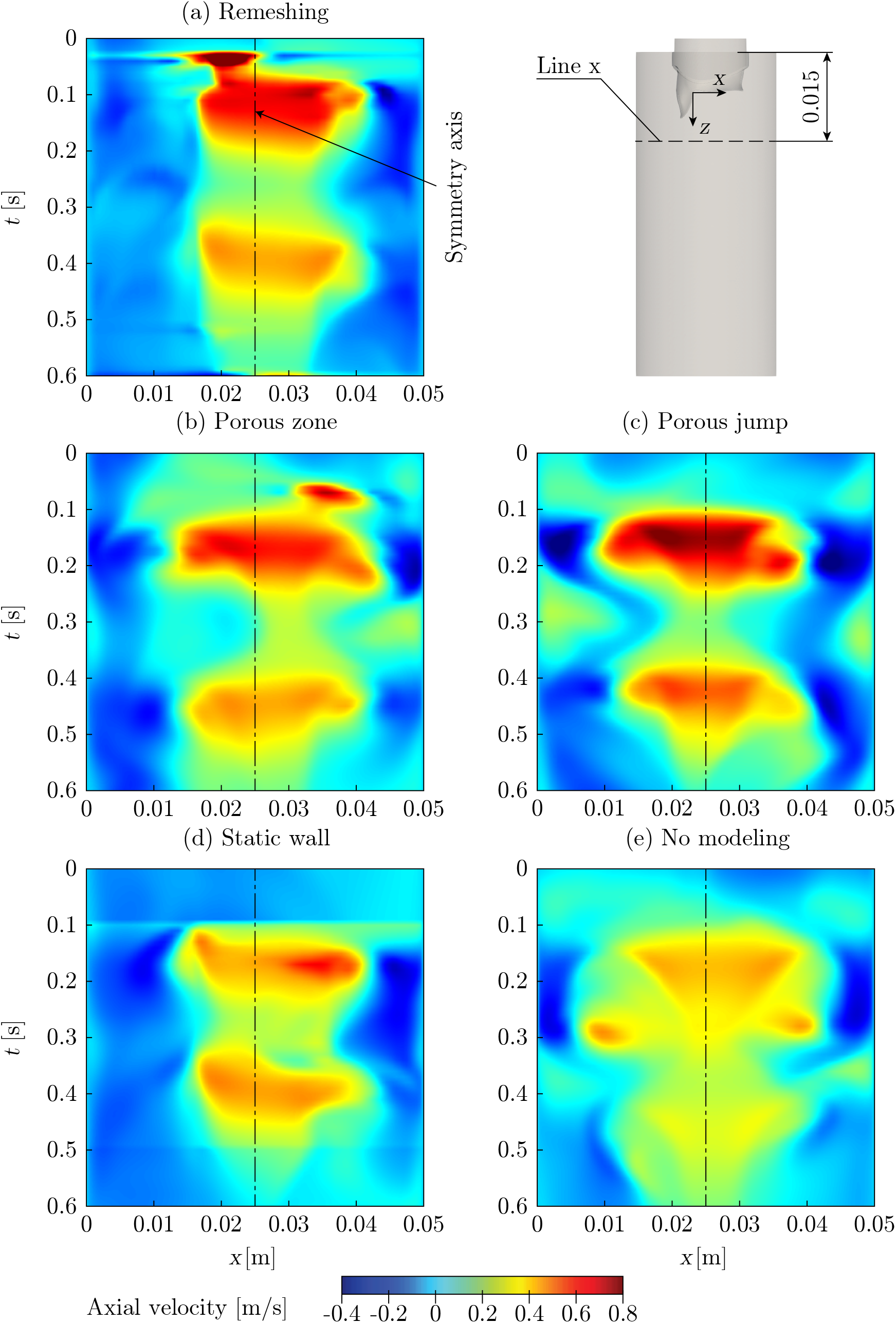
Spatio-temporal axial velocity along the x-direction below the MV. Dimensions in [m].

A global metric *S* for the quantification on asymmetry was determined according to equation 7. The velocity distribution for the model using remeshing results in *S* = 0.27. A similar level of asymmetry is obtained by the porous zone model, *S* = 0.3, and the model with a porous jump condition,*S* = 0.25. The most asymmetric flow with *S* = 0.37 is present in the simulation where the MV is modeled as a static wall. In contrast, the simulation without any MV yields *S* = 0.19.

The visualization of the vortex structures already revealed that the propagation velocity of the incoming mitral jet changes with the different modeling approaches. To quantify this, the temporal change of the velocity component in *z*-direction was assessed along the MA center line, which is indicated as *line-z* in figure 10. The resulting spatio-temporal profiles clearly show that the velocity wave generated by the velocity profile at the inlet propagates into the fluid domain. Its propagation speed is determined by tracking the peak velocity generated by the E-wave, whereas the delayed A-wave front shows a similar pattern but with a reduced axial velocity. The distance traveled is measured from the location of the MA. The resulting slope is linear and changes as the transmitral flow velocity changes from acceleration to deceleration, which can be observed for the E-wave in figure 2 at *t* ≈ 0.1 s. The simulation with a dynamic mesh yields the highest propagation speed with 1.67 m s and 0.22 m s in the acceleration and deceleration phases, respectively. The former reduces by about 40% and 34% to 1/m s and 1.11/m s, respectively, for the simulation with a porous zone and a porous jump. In the deceleration phase, however, the slope of both is comparable to that of the one-way FSI.

**FIGURE 10.**
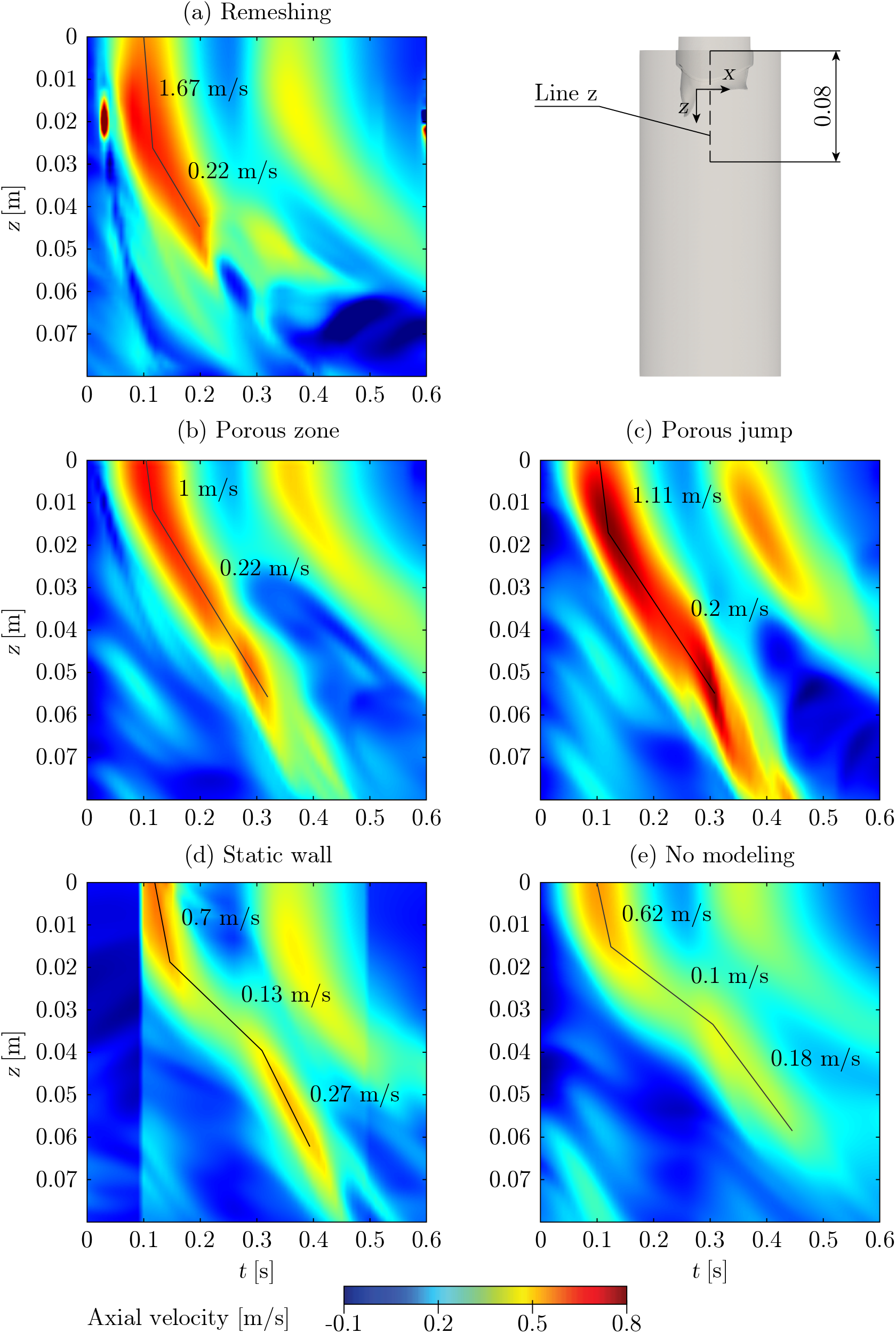
Spatio-temporal axial velocity along the MA center line. Black lines indicate the trajectory of the peak value of the velocity waves. Dimensions in [m].

Comparing both the model where the MV is implemented as a static wall and the model without any representation of the MV with the one-way FSI simulation, reveals that the propagation velocity is generally lower. In particular, in the acceleration phase, the slope decreases by about 60%, resulting in a smaller penetration depth of the main vortex (see figure 8 (b) and (c)). During the deceleration, the propagation velocity is slowed down by more than 40% and 50%, respectively. Moreover, their slopes change again at about *t* = 0.3 s to 0.27 m s and 0.18 m s, respectively.

### 3.4 Numerical costs and result summary

To evaluate the performance of each modeling approach, the wall-clock time per cardiac cycle was determined. All simulations were performed on an Intel Xeon CPU E5-2640 v4 with 2.4 GHz and 20 physical cores. The one-way FSI model was run on eight cores for the structural and 18 for the fluid simulation. All the other approaches were modeled using 20 cores. As the most expensive approach, the dynamic mesh model required about 14.5 h to solve one cardiac cycle. This was followed by the porous jump model with 4.2 h and the simulations in which the MV was modeled as a porous zone (1.6 h) and static wall (1.1 h), respectively. The shortest simulation time with *t* = 0.7 h was obtained by the simplest approach without any modeling of the MV.

A summary of the deviations for the various modeling approaches is shown in table 1 for the following analyzed values: maximum axial velocity *u*_*max*_ and phase-shift Δ*t*, cycle-averaged kinetic energy *w*_*kin*_, averaged transmitral pressure drop, flow asymmetry *S*, jet propagation speed *v*_*jet*_, and the numerical costs.

**TABLE 1.**
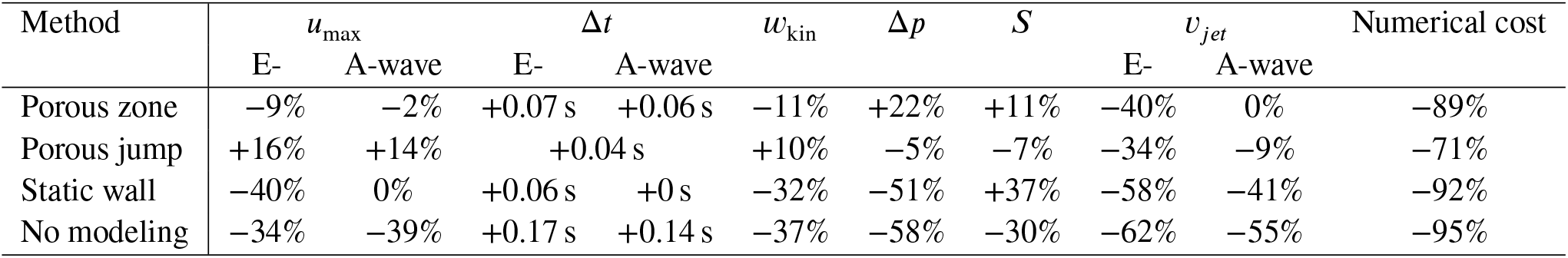
Deviations of the different approaches to the one-way FSI simulation. *u*_*max*_ - maximum axial velocity, Δ*t* - phase-shift, *w*_*kin*_ - cycle-averaged kinetic energy, - averaged transmitral pressure drop, *S* - flow asymmetry, *v*_*jet*_ - jet propagation speed

## 4 DISCUSSION

The aim of the work was to compare different modeling approaches of the MV in a consistent benchmark setup and provide quantitative metrics on the most important flow variables relevant for investigations of intraventricular flow patterns. A oneway FSI setup with remeshing was taken as the ground truth data and four other implementation approaches (porous zone, porous jump, static wall and no modeling) were compared to this setup. The capabilities of the ANSYS solver to perform FSI simulations were verified by reproducing a heart valve benchmark from literature. Velocity, kinetic energy, transmitral pressure drop, vortex formation, and flow asymmetry were investigated with qualitative and quantitative metrics.

Focussing on the velocities, all modeling approaches showed a phase shift of the peak E- and A-wave of up to 0.1 s, which corresponds to the time for opening and closing of a physiological human heart valve. In general, the orifice-like planar model of the MV can mimic the localized high transmitral velocities. However, because the 3D representation of the valve is missing, the cross section of the flow expands earlier leading to a flow field with reduced velocities behind the MV and a lower propagation speed. This effect is particularly strong in the simulation without any MV, which essentially models the flow through a tube with the diameter of the MA. Here, the peak velocity at the measured location is reduced by almost 40% and the propagation speed of the transmitral jet is lowered by more than 60%. This causes the phase shift to be greatest here. The maximum axial velocity for the porous jump model is increased by about 15%, compared to the one-way FSI simulation. Here, the reduced propagation speed and the resulting time delay of the E- and A-wave may be due to the time-dependent function of the resistive porous face.

The results of the one-way FSI simulation show a sudden drop in the kinetic energy per unit volume, which can be explained by the rapid opening of the MV, as the increasing opening area leads to a decreased transmitral flow velocity. The approaches that implement the MV as a static wall or do not a valve at all fail to accurately predict the energy density of the flow, especially in the opening phase of the MV. Moreover, the jump in the energy density with the static wall implementation indicates that this approach is very prone to numerical instabilities. The porous zone model matches the dynamic mesh model results quite well, although the energy density is underestimated during the E-wave when the MV opens and the resistance of the porous faces decreases. The opposite effect is observed for the model with porous jump condition. Here, the pressure drop is about 5% lower than in the dynamic remeshing model, while the average of the energy density is about 10% higher.

The presence of an MV leads to the formation of vortex structures at the tips of the valve leaflets instead of the MA if no MV is implemented, which was already found by ^13, 17^ and ^35^. The latter results in a more central position of the vortices and thus fails to produce a flow that is directed towards the septum. As a result, the vortex structures are less stable and start to disintegrate earlier, which was already observed by Seo et al. ^17^. In contrast to modeling the MV as a porous medium, where vortex formation starts in the MA plane, the vortex structures form further downstream in the tube of the test setup with the one-way FSI approach. The rolling-up process can already be observed in the opening phase, which cannot be captured with any other approach.

The approaches with porous jump condition and without MV produce a rather symmetric flow aligned with the central axis of the flow domain. The FSI, porous zone, and static wall approaches, however, are capable of deflecting the flow towards the lateral wall, which can be quantitatively confirmed by figure 9. This leads to an earlier disintegration of the vortex ring near the lateral wall, which is qualitatively shown in figure 7 and 8. These observations were also found in other numerical studies like ^17, 25^ and ^31^ and in clinical data of Charonko et al. ^35^.

As a common simplification, a computationally less expensive laminar solver was used because the flow in the left ventricle is mostly laminar and experiences turbulent transitions only locally and for short times. This is in agreement with other numerical studies like ^17, 30, 13^ and ^31^. Moreover, this particularly avoids problems with deforming inflation layers in dynamic mesh simulations. However, the cycle dependence of the simulation results indicates the presence of local turbulent effects and thus motivates the analysis of the influence of turbulence modeling.

The FSI simulation enables to mimic the opening and closing behavior of the MV, having an MV orifice area of about 4.6 cm^2^ and an opening angle of the anterior and posterior leaflet of about 90°. Both are within the physiological range reported in literature ^36^. However, there are several limitations in the modeling of the MV. The motion of the valve was defined by different loads on the leaflet surfaces, thus no two-way FSI simulation was performed, which would lead to a different behavior in the opening and closing phase. Moreover, there is no fluttering of the leaflets during the diastole, which was observed in the numerical studies of Govindarajan et al. ^31^ and Lee et al. ^37^. Instead, the MV opens continuously until it is fully open at *t* = 0.1 s. The leaflets were modeled as hyperelastic materials with isotropic properties that are generally in the physiological range. However, when extending the FSI simulation to a two-way coupling, a nonlinear and anisotropic behavior might be considered. In addition, chordae tendineae could be included to define the end-diastolic position of the anterior and posterior leaflets. However, the complexity and uncertainty of structural modeling makes the computational effort questionable, especially since a two-way coupling does not necessarily imply higher modeling fidelity with respect to hemodynamics ^38, 39^ and ^40^. As in the study of Canè et al., no movement of the MA was considered for simplicity ^13^. In contrast, the atrium is contracting in the model presented by Daub, resulting in a motion of the MV plane with time and an additional but weaker transmitral jet entering the ventricle ^30^.

The one-way FSI simulation takes about nine times longer to simulate a cardiac cycle than all other approaches, which is mainly due to the spatial discretization with an unstructured grid. Dynamic remeshing is not compatible with hanging nodes or polyhedral elements, thus tetrahedral elements had to be used. For this reason, the mesh size had to be increased by a factor of four. In addition, applying mesh smoothing and remeshing at each time step increases the numerical cost compared to all other approaches.

Regarding the patient-specific implementation of the MV geometry, the user either needs a time-dependent projection of the effective opening area or a full 3D reconstruction of the MV. The latter might be solved by recent approaches that sucessfully segmented a MV using transesophageal echocardiography ^41^, computed tomography ^42^ or magnetic resonance imaging ^43^. Despite these solutions, MV segmentation is not a trivial task and the porosity approach might be advantageous for specific research questions considering its superiority regarding flexibility and ease-of-implementation.

## 5 CONCLUSION

The presented framework compared different modeling approaches of the MV. The rather qualitative assessments found in the literature have been extended by a set of quantitative metrics. The porous medium approach already presented in the literature was extended by a time-dependent function for porosity, which was found to be very important for mimicking the flow and predicting flow features such as pressure drop and kinetic energy. A further simplification to a planar 2D model did not improve numerical stability or reduce computation time. The benchmark setup for reproduction of the results and testing of other MV modeling approaches is available under 10.5281/zenodo.6741579.

In summary, the findings of this work make it possible to formulate general guidelines for the various modeling strategies with respect to different flow variables and the complexity of implementation, which are listed in table 2.

**TABLE 2.** Guidelines for MV modeling. FSI - One-way fluid structure interaction with remeshing, PZ - porous zone, PJ - porous jump, SW - static wall, NO - no modeling.

| Method | FSI | PZ | PJ | SW | NO |
| --- | --- | --- | --- | --- | --- |
| Kinetic energy | + | + | + | – | – |
| Pressure drop | + | + | + | – | – |
| Flow asymmetry | + | + | – | + | – |
| Jet propagation speed | + | + | + | – | – |
| Numerical cost | – | + | + | + | + |
| Vortices | + | + | – | – | – |
| Turbulence | – | + | + | + | + |
| Implementation | – | + | + | – | + |

## Ethics statement

None required.

