## Supplementary Information for "Generic Framework for Quantifying the Influence of the Mitral Valve on Ventricular Blood Flow"

#### Validation of fluid-structure interaction algorithm

The displacement of the flap in $x$- and $y$-direction for the overset and remeshing approaches are shown in figure 1. Both segregated and coupled pressure-velocity coupling schemes were tested.

The deformation of the flap increases as the velocity at the inflow increases and reaches its maximum at $t \approx 0.3\,\mathrm{s}$. After that the inlet velocity decreases causing the flap to rebound until one cycle is completed at $t \approx 0.8\,\mathrm{s}$. Given the good agreement with findings from [1] the FSI algorithm can be approved.

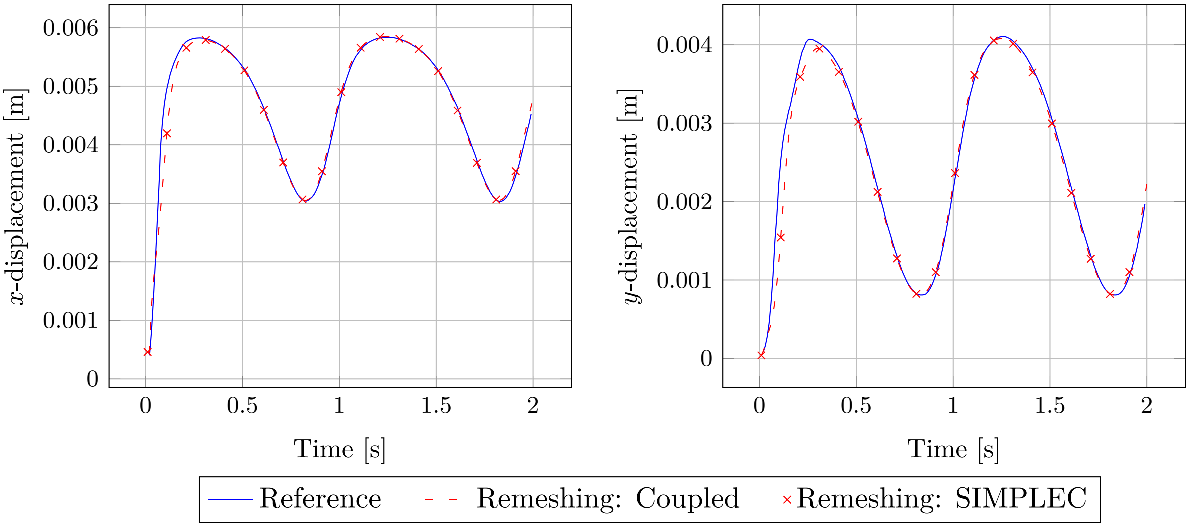


**Figure 1:** Displacements in x- and y-direction of the bottom leaflet tip, computed with different methods. Reference taken from [1].

In figure 2 the flow field of the domain is visualized using the LIC technique with the color denoting the velocity magnitude. In this method, a white noise is first applied to the fluid region. Then, for each pixel in that image, all neighboring pixels are identified that lie on the forward and backward streamlines with fixed arc length. The gray level of the current pixel is then calculated by a texture convolution, with weights determined by a low-pass filter applied to the previously identified set of pixels. This finally results in a gray level LIC image with points on the same field line having a much greater similarity than points on adjacent field lines [2]. The constriction caused by the flap results in an increased local fluid velocity. A critical point can be identified behind the flap which marks a region where the magnitude of the velocity is zero and where recirculation of the flow is present.

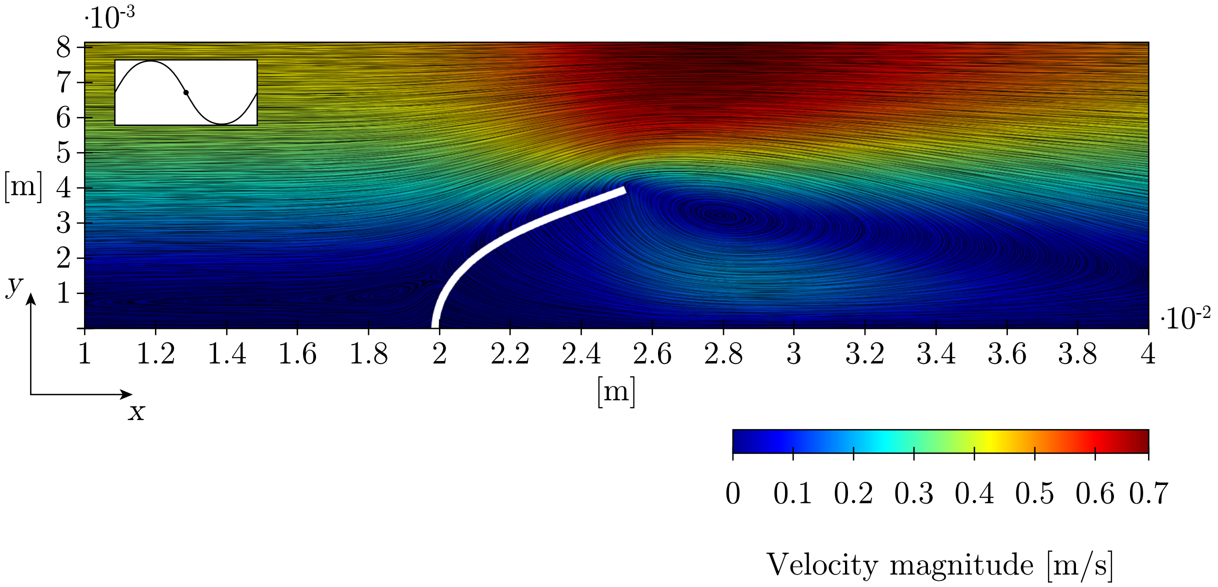


**Figure 2:** LIC visualization superimposed on a velocity magnitude contour plot, at $t = 0.5\,\mathrm{s}$ of the simulation using remeshing and smoothing only.

#### Validation of porous medium model

To evaluate the quality of the results obtained by all approaches using porous media models, the pressure drop over the channel length is reported in figure 3. The pressure drop increases as the velocity at the inlet increases. Moreover, the numerical results match perfectly with the analytical solution.

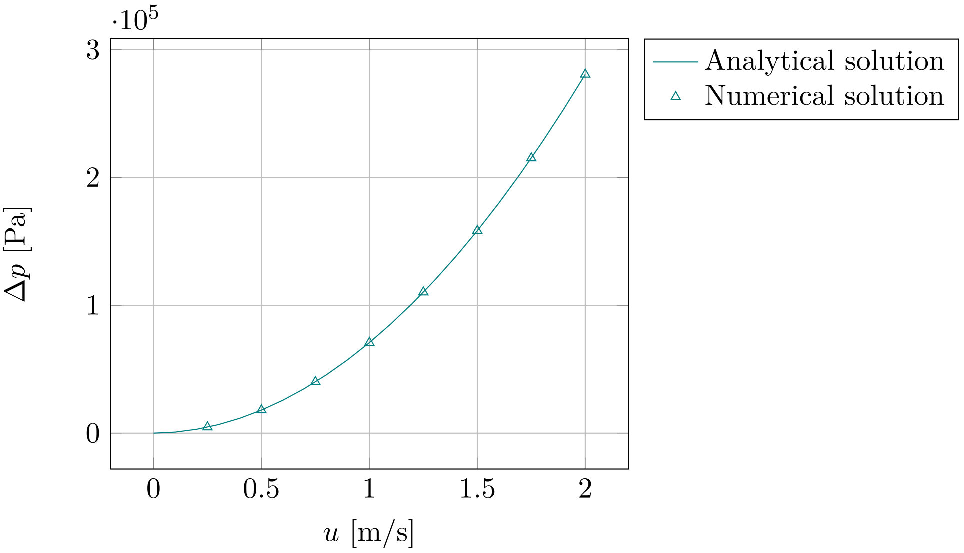


**Figure 3:** Pressure drop over the channel length for different flow velocities at the inlet.

#### Loads on the mitral valve surfaces for one-way FSI

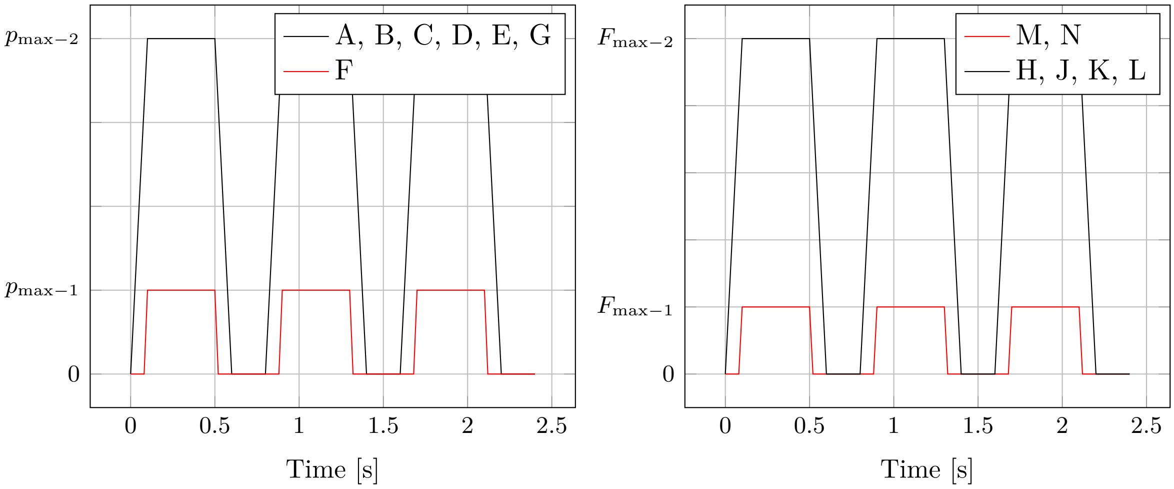


**Figure 4:** Left: pressure load on MV leaflets. Right: forces on edges of MV geometry.

**Table 1:** Maximum loads on different regions of the MV.

| Label | A | B | C | D | E | F | G | H | J | K | L | M | N |
|---|---|---|---|---|---|---|---|---|---|---|---|---|---|
| Max. pressure [MPa] | 20 | 15 | 10 | 15 | 17.5 | 5 | 15 | - | - | - | - | - | - |
| Max. force [N] | - | - | - | - | - | - | - | 0.05 | 0.2 | 0.15 | 0.1 | 0.075 | 1 |

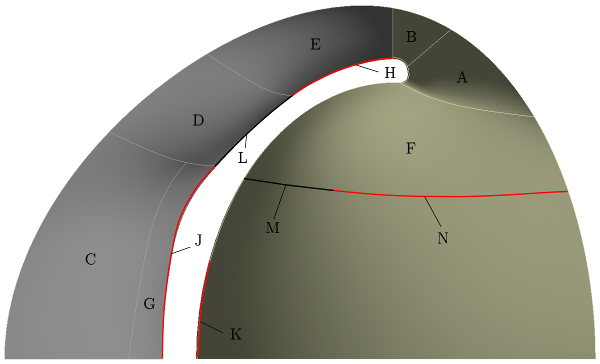


**Figure 5:** Labels of MV geometry having different loads.

#### Mesh independence study

To estimate the discretization error, variables must be selected that are important to the objective of the following simulation studies. Therefore, the mean value over the whole simulation time of the area-averaged velocity magnitude at the symmetry plane and the integral of the static pressure at the FSI interface are reported. The results are shown in figure 6. The results are very similar for all grid resolutions. Since the following analyses will focus mainly on local effects, the rather fine mesh with about 1.38 million cells will be used in all subsequent simulations.

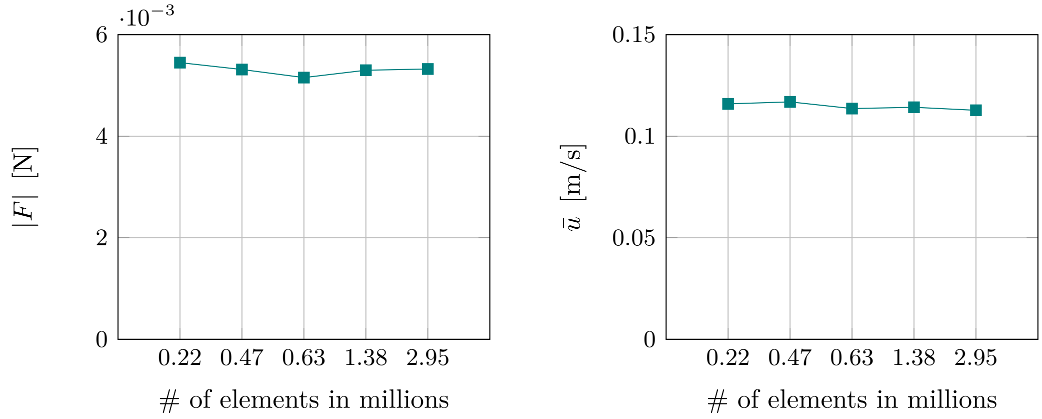


**Figure 6:** Left: absolute value of integral of static pressure at the FSI interface. Right: time-average of area-averaged velocity magnitude at symmetry plane.

### Porous medium setup

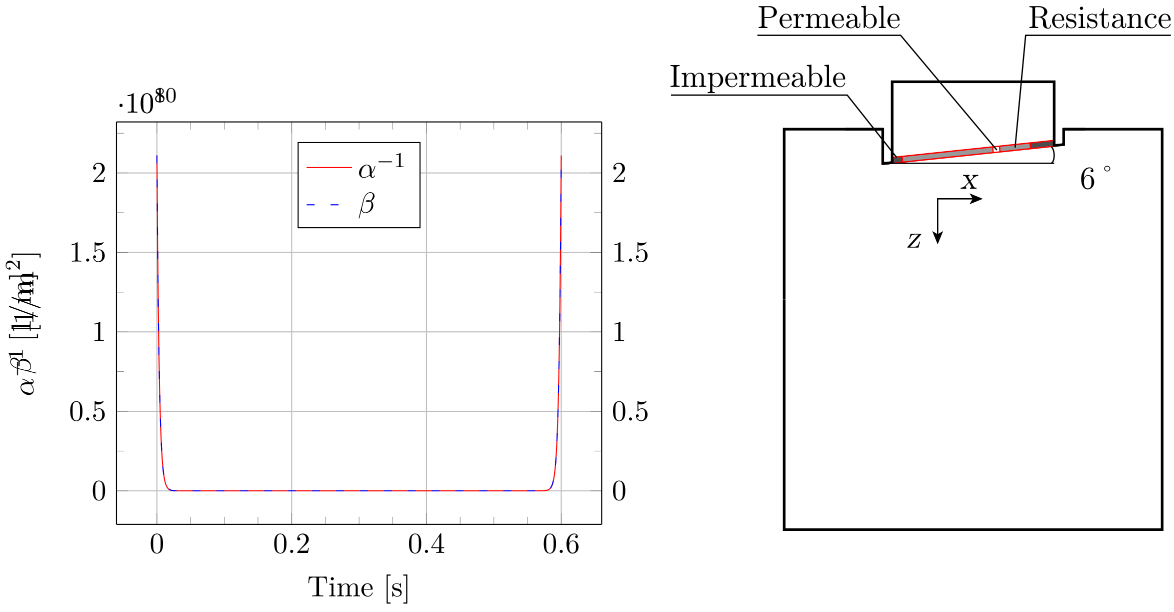


**Figure 7:** Left: inverse permeability $\alpha^{-1}$ and Forchheimer coefficient $\beta$ as a function of time for the diastole. Right: projected MV geometry in the benchmark setup.

### Porous media properties in Ansys Fluent

This section collects the expressions implemented for the porous zone model and the simulation with porous jump condition.

#### Porous zone

The inverse permeability $\alpha^{-1}$, which is the *viscous resistance* in Fluent, in the normal direction of the MV plane was defined as follows

```
IF(AND(t >= 2.1[s], t < 2.2[s]), 2.11e+8[m^-1]*exp((t-2.2[s])*300[s^-1]),
IF(AND(t >= 1.7[s], t < 2.1[s]), 0[m^-1],
IF(AND(t >= 1.6[s], t < 1.7[s]), 2.11e+8[m^-1]*exp(-(t-1.6[s])*300[s^-1]),
IF(AND(t >= 1.4[s], t < 1.6[s]), 2.11E+8[m^-1],
IF(AND(t >= 1.3[s], t < 1.4[s]), 2.11e+8[m^-1]*exp((t-1.4[s])*300[s^-1]),
IF(AND(t >= 0.9[s], t < 1.3[s]), 0[m^-1],
IF(AND(t >= 0.8[s], t < 0.9[s]), 2.11e+8[m^-1]*exp(-(t-0.8[s])*300[s^-1]),
IF(AND(t >= 0.6[s], t < 0.8[s]), 2.11E+8[m^-1],
IF(AND(t >= 0.5[s], t < 0.6[s]), 2.11e+8[m^-1]*exp((t-0.6[s])*300[s^-1]),
IF(AND(t >= 0.1[s], t < 0.5[s]), 0[m^-1],
IF(t < 0.1[s], 2.11e+8[m^-1]*exp(-t*300[s^-1]), 2.11E+8[m^-1]))))))))))
```

In contrast, all other directions in which the flow is blocked were defined as follows

```
IF(AND(t >= 2.1[s], t < 2.2[s]), 2.11e+8[m^-1]*exp((t-2.2[s])*300[s^-1])
    +211000[m^-1],
IF(AND(t >= 1.7[s], t < 2.1[s]), 211000[m^-1],
IF(AND(t >= 1.6[s], t < 1.7[s]), 2.11e+8[m^-1]*exp(-(t-1.6[s])*300[s^-1])
    +211000[m^-1],
IF(AND(t >= 1.4[s], t < 1.6[s]), 2.11E+8[m^-1],
IF(AND(t >= 1.3[s], t < 1.4[s]), 2.11e+8[m^-1]*exp((t-1.4[s])*300[s^-1])
    +211000[m^-1],
```

```
IF(AND(t >= 0.9[s], t < 1.3[s]), 211000[m^-1],
IF(AND(t >= 0.8[s], t < 0.9[s]), 2.11e+8[m^-1]*exp(-(t-0.8[s])*300[s^-1])
    +211000[m^-1],
```

The Forchheimer coefficient $\beta$, which is the *inertial resistance* $C_2$ in Fluent, in the primary flow direction was defined as follows

```
IF(AND(t >= 2.1[s], t < 2.2[s]), 2.11e+10[m^-2]*exp((t-2.2[s])*300[s^-1])
    +1[m^-2],
IF(AND(t >= 1.7[s], t < 2.1[s]), 1[m^-2],
IF(AND(t >= 1.6[s], t < 1.7[s]), 2.11e+10[m^-2]*exp(-(t-1.6[s])*300[s^-1])
    +1[m^-2],
IF(AND(t >= 1.4[s], t < 1.6[s]), 2.11e+10[m^-2],
IF(AND(t >= 1.3[s], t < 1.4[s]), 2.11e+10[m^-2]*exp((t-1.4[s])*300[s^-1])
    +1[m^-2],
IF(AND(t >= 0.9[s], t < 1.3[s]), 1[m^-2],
IF(AND(t >= 0.8[s], t < 0.9[s]), 2.11e+10[m^-2]*exp(-(t-0.8[s])*300[s^-1])
    +1[m^-2],
IF(AND(t >= 0.6[s], t < 0.8[s]), 2.11e+10[m^-2],
IF(AND(t >= 0.5[s], t < 0.6[s]), 2.11e+10[m^-2]*exp((t-0.6[s])*300[s^-1])
    +1[m^-2], IF(AND(t >= 0.1[s], t < 0.5[s]), 1[m^-2],
IF(t < 0.1[s], 2.11e+10[m^-2]*exp(-t*300[s^-1])
    +1[m^-2], 2.11e+10[m^-2]))))))))))
```

In the directions which behave like a solid wall, the expression for $\beta$ reads

```
IF(AND(t >= 2.1[s], t < 2.2[s]), 2.11e+10[m^-2]*exp((t-2.2[s])*300[s^-1])
    +211000[m^-2],
IF(AND(t >= 1.7[s], t < 2.1[s]), 211000[m^-2],
IF(AND(t >= 1.6[s], t < 1.7[s]), 2.11e+10[m^-2]*exp(-(t-1.6[s])*300[s^-1])
    +211000[m^-2],
IF(AND(t >= 1.4[s], t < 1.6[s]), 2.11E+10[m^-2],
IF(AND(t >= 1.3[s], t < 1.4[s]), 2.11e+10[m^-2]*exp((t-1.4[s])*300[s^-1])
    +211000[m^-2],
IF(AND(t >= 0.9[s], t < 1.3[s]), 211000[m^-2],
IF(AND(t >= 0.8[s], t < 0.9[s]), 2.11e+10[m^-2]*exp(-(t-0.8[s])*300[s^-1])
    +211000[m^-2],
IF(AND(t >= 0.6[s], t < 0.8[s]), 2.11E+10[m^-2],
IF(AND(t >= 0.5[s], t < 0.6[s]), 2.11e+10[m^-2]*exp((t-0.6[s])*300[s^-1])
    +211000[m^-2],
IF(AND(t >= 0.1[s], t < 0.5[s]), 211000[m^-2],
IF(t < 0.1[s], 2.11e+10[m^-2]*exp(-t*300[s^-1])
    +211000[m^-2], 2.11E+10[m^-2]))))))))))
```

#### Porous jump

The face permeability $\alpha$ was defined as follows

```
IF(AND(t >= 2.1[s], t < 2.2[s]), 1e-09[m^2]*exp(-(t-2.2[s])*207.23[s^-1]),
IF(AND(t >= 1.7[s], t < 2.1[s]), 1[m^2],
IF(AND(t >= 1.6[s], t < 1.7[s]), 1e-09[m^2]*exp((t-1.6[s])*207.23[s^-1]),
IF(AND(t >= 1.3[s], t < 1.4[s]), 1e-09[m^2]*exp(-(t-1.4[s])*207.23[s^-1]),
IF(AND(t >= 0.9[s], t < 1.3[s]), 1[m^2],
IF(AND(t >= 0.8[s], t < 0.9[s]), 1e-09[m^2]*exp((t-0.8[s])*207.23[s^-1]),
IF(AND(t >= 0.5[s], t < 0.6[s]), 1e-09[m^2]*exp(-(t-0.6[s])*207.23[s^-1]),
IF(AND(t >= 0.1[s], t < 0.5[s]), 1[m^2],
IF(t < 0.1[s], 1e-09[m^2]*exp(t*207.23[s^-1]), 1e-09[m^2]))))))))
```

The Forchheimer coefficient $\beta$, which is the *pressure-jump coefficient* $C_2$ in Fluent, was defined as follows

```
IF(AND(t >= 2.1[s], t < 2.2[s]), 1e+06[m^-1]*exp((t-2.2[s])*250[s^-1]),
IF(AND(t >= 1.7[s], t < 2.1[s]), 0[m^-1],
IF(AND(t >= 1.6[s], t < 1.7[s]), 1e+06[m^-1]*exp(-(t-1.6[s])*250[s^-1]),
IF(AND(t >= 1.3[s], t < 1.4[s]), 1e+06[m^-1]*exp((t-1.4[s])*250[s^-1]),
IF(AND(t >= 0.9[s], t < 1.3[s]), 0[m^-1],
IF(AND(t >= 0.8[s], t < 0.9[s]), 1e+06[m^-1]*exp(-(t-0.8[s])*250[s^-1]),
IF(AND(t >= 0.5[s], t < 0.6[s]), 1e+06[m^-1]*exp((t-0.6[s])*250[s^-1]),
IF(AND(t >= 0.1[s], t < 0.5[s]), 0[m^-1],
IF(t < 0.1[s], 1e+06[m^-1]*exp(-t*250[s^-1]), 1e+06 [m^-1])))))))))
```

#### Static wall setup

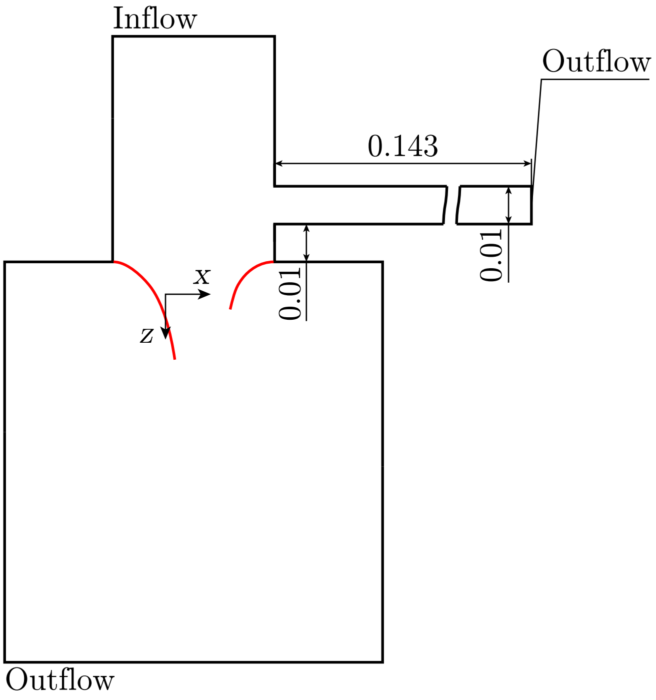


**Figure 8:** Adaptation of the benchmark setup for the approach modeling the MV as a static wall

##### Flow rate weighting for static wall approach

The fractional flow rate through main exit was defined as

```
IF(AND(t >= 1.7[s], t < 2.1[s]), 1,
IF(AND(t >= 0.9[s], t < 1.3[s]), 1,
IF(AND(t >= 0.1[s], t < 0.5[s]), 1, 0)))
```

In contrast, the *flow rate weighting* for the outflow above the MV reads

```
IF(AND(t >= 1.7[s], t < 2.1[s]), 0,
IF(AND(t >= 0.9[s], t < 1.3[s]), 0,
IF(AND(t >= 0.1[s], t < 0.5[s]), 0, 1)))
```
